# BCG overexpressing an endogenous STING agonist provides enhanced protection against pulmonary tuberculosis

**DOI:** 10.1101/365411

**Authors:** Ruchi Jain Dey, Bappaditya Dey, Alok Kumar Singh, Monali Praharaj, William Bishai

## Abstract

Stimulator of interferon genes (STING) has emerged as a key signaling receptor that induces proinflammatory cytokines, and small molecule STING agonists are being developed as anticancer and antiviral agents. Here we report a strategy of delivering a STING agonist from within live BCG. We generated a recombinant BCG (BCG-*disA*-OE) that overexpresses the endogenous mycobacterial diadenylate cyclase gene and releases high levels of the STING agonist c-di-AMP. In macrophages BCG-*disA*-OE elicited statistically significantly stronger TNF-α, IL-6, IL-1β, IRF3, and IFN-β levels than BCG-WT. In a 24-week guinea pig vaccination-*Mtb* challenge model, BCG-*disA*-OE reduced lung weights, pathology scores, and *Mtb* CFU counts in lungs by 28% (p<0.05), 34%, and 2.0 log10 CFU units (p < 0.5) compared with BCG-WT, respectively. Overproduction of the STING agonist c-di-AMP significantly enhanced the protective efficacy of BCG against pulmonary and extrapulmonary tuberculosis. Our findings support the development of BCG-vectored STING agonists as a TB vaccine strategy.

## INTRODUCTION

Recent studies have identified a key role for the stimulator of interferon genes (STING) intracellular sensor in mediating innate immune responses cellular stress or pathogen infection [1,2]. STING is a cytosolic receptor for both pathogen-associated molecular pattern (PAMP) molecules such as cyclic dinucleotides c-di-AMP or c-di-GMP produced by bacteria, and for mammalian endogenous danger signaling DAMP molecules such as 2’,3’ cyclic GMP-AMP (cGAMP) which is synthesized by cGAS (cyclic GMP-AMP synthase) in response to microbial or self-derived cytosolic double-stranded DNA [1-4]. Activation of STING induces numerous interferon-stimulated genes including type I interferons (IFNα/β) and is associated with co-activation of NF-κB and STAT6 transcription factors that follows or parallels the induction of interferon regulatory factors (IRF) transcription factors. Thus endogenous and exogenous cyclic dinucleotides (CDNs) are strong TLR-independent mediators of innate host defenses [5,6]. Based on the key role of STING signaling, pharmacological stimulation of the pathway using small molecule STING agonists is currently being tested for enhancement of antitumor immunity and as potential anti-viral therapy [5,7].

As they are capable of inducing potent cytokine and cellular immune responses against pathogens, CDN STING agonists also exhibit attractive vaccine adjuvant properties. They increase expression of MHC class II, co-stimulatory molecules (CD80/CD86) as well as activation (CD40) and adhesion (CD45) markers; in addition STING agonists have been shown to enhance antigen processing and presentation to T cells [8-10]. Among microbial-derived CDNs, c-di-AMP has emerged as an efficient activator of macrophages and leads to robust Th1, Th17 and CD8 T cell responses [11].

Efforts to harness small molecule CDNs themselves as vaccine adjuvants to enhance systemic immunity, however, may be limited by rapid systemic clearance and by the fact that as negatively charged molecules, CDNs do not effectively cross cell membranes of macrophages and professional antigen presenting cells [6]. Hence there is a need to develop a cost-effective formulations of CDNs and related STING agonists that allow for sustained release to the cytosolic compartment of antigen-presenting cells.

Our previous findings showed that *Mycobacterium tuberculosis* (*Mtb*), an intracellular pathogen, possesses di-adenylate cyclase enzyme, *disA* (MT3692), that synthesizes and secretes c-di-AMP into the host cell cytosol. We showed that an *Mtb* strain engineered to overexpress c-di-AMP (*Mtb-disA*-OE) was significantly attenuated in lethality, ability to proliferate, and cause disease pathology in a mouse model [12]. In a separate study, we showed that mutation of the *cdnP* gene encoding a cyclic dinucleotide phosphodiesterase (CdnP) that hydrolyzes c-di-AMP lead to a mutant Mtb strain that accumulates c-di-AMP and is similarly attenuated for virulence [13].

Based on these results we constructed a c-di-AMP–overexpressing bacillus Calmette–Guérin (BCG), an attenuated strain of *Mycobacterium bovis* widely used as a TB vaccine globally, and found that it induced a significantly higher IRF and IFN-β response than BCG itself, indicating that bacterial-derived c-di-AMP gains access to the host cell cytosol despite the fact that BCG lacks the ESX-1 protein secretion system found in *Mtb*. These findings strongly encouraged us to evaluate whether BCG-induced immune responses against TB are boosted by overexpression of the STING agonist c-di-AMP.

A number of candidate vaccines for TB are currently being evaluated in clinical trials, prime boost strategies that include BCG have failed to show improved protective efficacy in humans over BCG alone [14]. In the paper we report the development of a recombinant BCG which is boosted by endogenous overexpression of a STING agonist. This c-di-AMP–overexpressing recombinant BCG strain (BCG-*disA*-OE) exhibited increased IRF induction, IFN-β synthesis, and release of the pro-inflammatory cytokines IL-6, TNFα and IL1β in vitro than were observed with BCG-WT. Importantly, guinea pigs vaccinated with BCG-*disA*-OE were significantly better protected against aerosol challenge with virulent *Mtb* than with BCG-WT suggesting improved protective efficacy over the existing BCG strain.

## Results

### Induction of type I IFN responses by BCG-*disA*-OE

We constructed a recombinant BCG strain (BCG-*disA*-OE) overexpressing the endogenous diadenylate cyclase gene *disA* (also known as *dacA*) encoded by MT3692 in the CDC1551 genome or Rv3568 in the H37Rv genome. The *disA* genes of *Mtb* and BCG are 100% identical at the nucleotide level. DisA catalyzes the conversion of 2 ATP molecules to c-di-AMP (**Supplementary Figure 1a**, **1b** and **1c**). The overexpression construct was generated by fusing the *disA* gene to the strong mycobacterial promoter *hsp60* within the episomal mycobacterial overexpression vector pSD5-hsp60. Gene expression profiling by real time PCR showed ~50-fold upregulation of *disA* expression in BCG-*disA*-OE as compared to the parental strain (**Supplementary Figure 1d**).

Next we evaluated whether strong *disA* expression would result in c-di-AMP-mediated IRF3 activation and consequent elevations in IFN-β levels in mouse macrophages. Raw Blue™ reporter macrophage cells when infected with BCG-*disA*-OE strains showed a significant 2-fold induction of IRF3 as compared to that observed with the wild-type parental strain BCG-Pasteur-WT (**Supplementary Figure 2a**). These preliminary results suggested that heightened levels of c-di-AMP release from BCG-*disA*-OE produced significant activation of the STING/IRF3 axis, an observation similar to our earlier findings with Mtb-*disA*-OE [12].

Next we infected primary murine BMDMs with BCG-*disA*-OE and BCG-WT and quantified *Ifnb* gene expression using qPCR. BCG-*disA*-OE-infected macrophages showed a 2-fold induction of *Ifnb* (p < 0.005, **Supplementary Fig 2b**) during an early temporal window. The notion that CDNs such as c-di-AMP can induce STING-dependent, but c-GAS independent induction of type I IFN responses (even in absence of extracellular DNA) was validated earlier, and these results were in accordance with our previous findings with recombinant *M.tb* overexpressing *disA* [12]. Previous studies suggest that viral infection of non-phagocytic cells lead to cytosolic penetration by leaking CDNs thus resulting into IFNβ production [18]. Since BCG lacks a functional Esx-1 secretion system required to release bacterial DNA, these results further reinforce the idea that phagosomes harboring mycobacteria are rather dynamic leaky structures or really do not require Esx-1 for membrane disruption [19]. Our data indicate that bacterial-derived c-di-AMP is detected in the macrophage cytosol and leads to STING-dependent IFN-β synthesis. Increased levels of type I IFNs in macrophages in response to increased c-di-AMP production in genetically modified BCG was the first step towards generation of a vaccine strain with an increased antigenic repertoire and ability to stimulate STING.

### Macrophage activation by c-di-AMP overexpressing BCG (BCG-*disA*-OE)

Although, the binding affinity of c-di-AMP for STING is weaker than that for cGAMP, ligation of c-di-AMP with STING appears sufficient to induce co-activation of transcription factors other than IRF-3 and, hence induction of pro-inflammatory cytokines [20,21]. Identification of another bona-fide physiological sensor for c-di-AMP, an endoplasmic adaptor, ERAdP, that binds to c-di-AMP with higher affinity, suggests ERAdP-dependent initiation of activation of NF-KB signaling in innate immune cells during bacterial infection [20]. We previously showed that *disA*-OE strains of *M.tb* induce a strong pro-inflammatory cytokines, such as TNF-α, IL-6 and IL-1β, suggesting macrophage activation and a complex link between c-di-AMP-based STING activation and induction of pro-inflammatory cytokines and other interferon stimulated genes (ISGs) [12]. Bone marrow-derived primary murine macrophages infected with BCG-*disA*-OE showed significant increased levels of TNF-α, IL-6 and IL-1β in culture supernatants as compared to uninfected or BCG-WT-infected controls (**Figure 1**). These results reveal a robust macrophage activation phenotype in response to c-di-AMP overproducing BCG with increased levels of M1 or Th1 cytokines (TNF-α, IL-1β and IL-6) that is more pronounced than that seen with BCG-WT.

**Figure 1.**
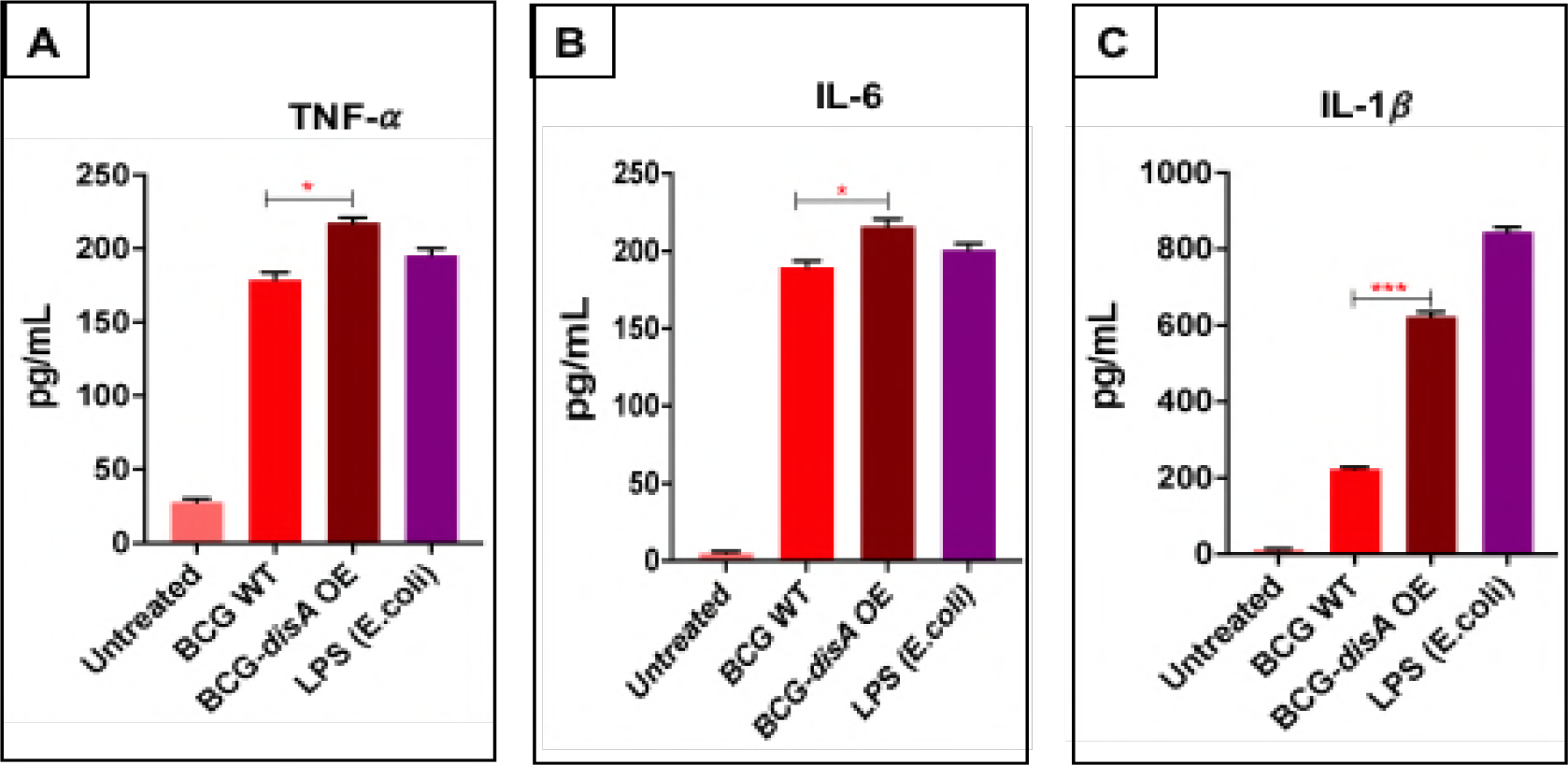
Modulation of pro-inflammatory cytokines in response to *disA* overexpression. Differential induction of **(a)** TNF-*α*, **(b)** IL-6 and **(c)** IL1*β* in mouse BMDMs challenged with wild-type and *disA* overexpression strains of *Mycobacterium bovis* BCG-Pasteur. BMDMs were challenged with wild-type and *disA* OE strains at an MOI of 1:20 for 5 h to establish the infection. Uninfected bacteria were washed using ice423 cold DPBS and cells were subsequently incubated for another 24 h. Culture supernatants were assayed by ELISA for different cytokines. The graphical points represent mean of 3 independent experiments ± standard error mean (SEM). Student’s t test (*P < 0.05 **P < 0.01, ***P < 0.001). MOI (multiplicity of infection).

### CDN-adjuvanted recombinant BCG offers better protection against virulent *M.tb* challenge in guinea pigs

Our *in vitro* studies accessing macrophage response due to increased levels of c-di-AMP encouraged us to test the vaccine potential of BCG-*disA*-OE in the guinea pig model of tuberculosis infection (**Supplementary Figure 3**). Groups of twelve guinea pigs were vaccinated intradermally with 0.1 ml of PBS (sham vaccination), 10^5^ CFU of BCG-*disA*-OE or BCG-WT and held for six weeks before challenge with aerosol challenge with 10^2^ CFU of *M. tuberculosis* H37Rv. As described in the Methods, lungs from one set of infected animals were obtained on day 1 after challenge to confirmed this implantation dose of *M. tuberculosis* (**Supplementary Figure 4**). Separate groups of infected animals were euthanized 14 and 18 weeks post-challenge to determine the protective efficacies of BCG-*disA*-OE and BCG-WT by organ weight, gross pathology, and bacterial loads in lungs and spleen.

At 14-week post-challenge, both BCG-WT and BCG-*disA-*OE vaccinated animals showed significantly lower lung weight, gross pathology scores, and the bacillary loads in the lungs, relative to saline-treated controls (**Figure 2A** and **2B**). Vaccination with BCG-*disA-*OE resulted in the highest reduction in lung gross pathology score, which was even more pronounced at 18-week post-challenge (**Figure 3A** and **3B**). While the impact of BCG-*disA*-OE vaccination on lung CFU counts was modest at the 14 week time point, by the 18 week time point lung CFU counts in the BCG-*disA*-OE vaccinated guinea pigs were 2.0 log10 units lower than in animals vaccinated with BCG-WT. In fact, two out of six guinea pigs in BCG-*disA*-OE group had lung CFU counts below the limit of detection which is ~3-5 bacilli.

**Figure 2.**
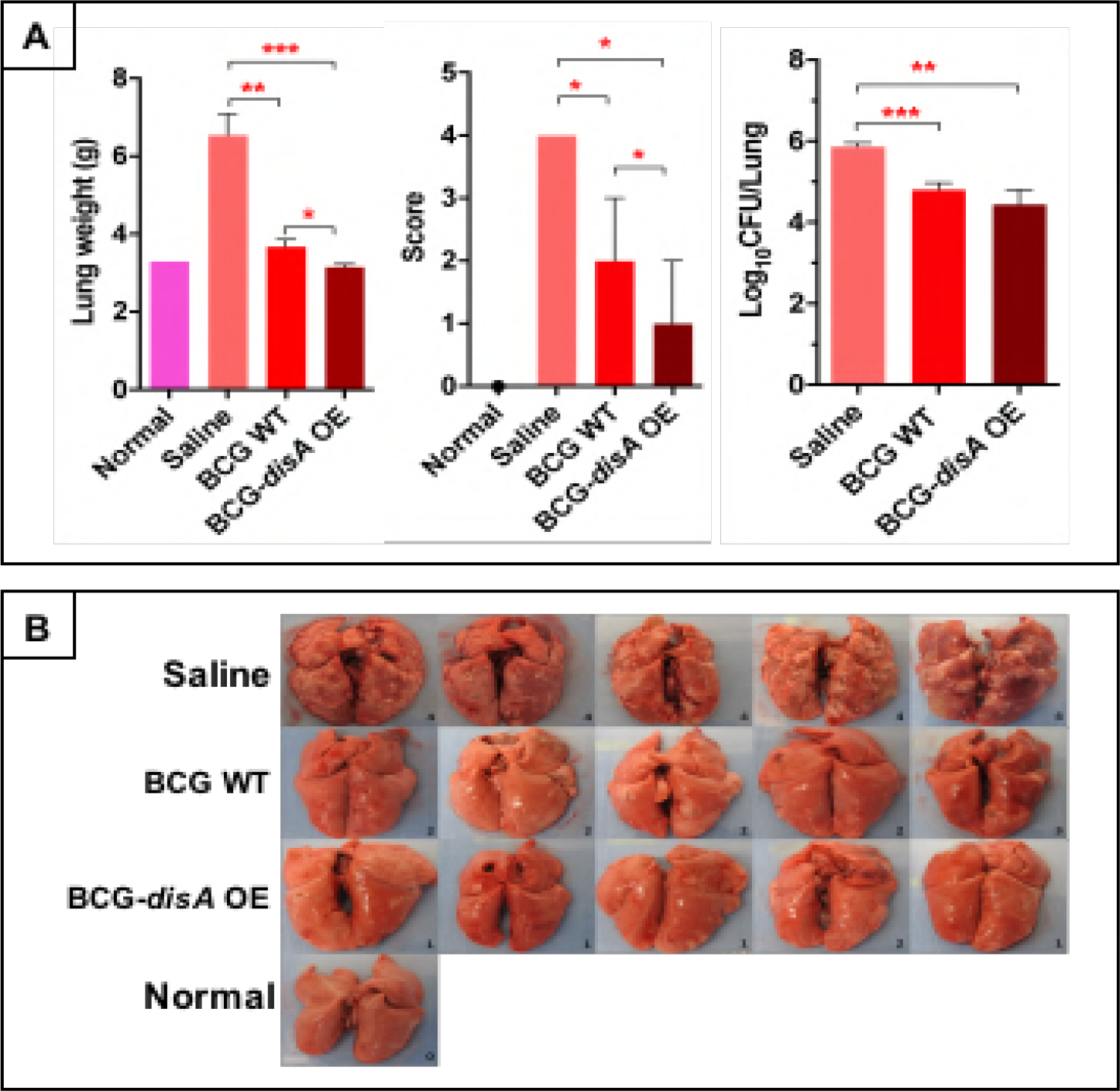
Effect on lung weight, gross lung pathology scores and lung gross 434 morphological features in guinea pigs 14 weeks post-challenge with *M. tuberculosis* H37Rv following vaccination with BCG-WT or BCG-*disA*-OE. A) Lung weights and gross pathology scores at 14 weeks post-*M. tuberculosis* challenge. B) Images of lungs at necropsy. ***P < 0.001, **, P< 0.01; *, P<0.05; Non-parametric Man Whitney Test.

**Figure 3.**
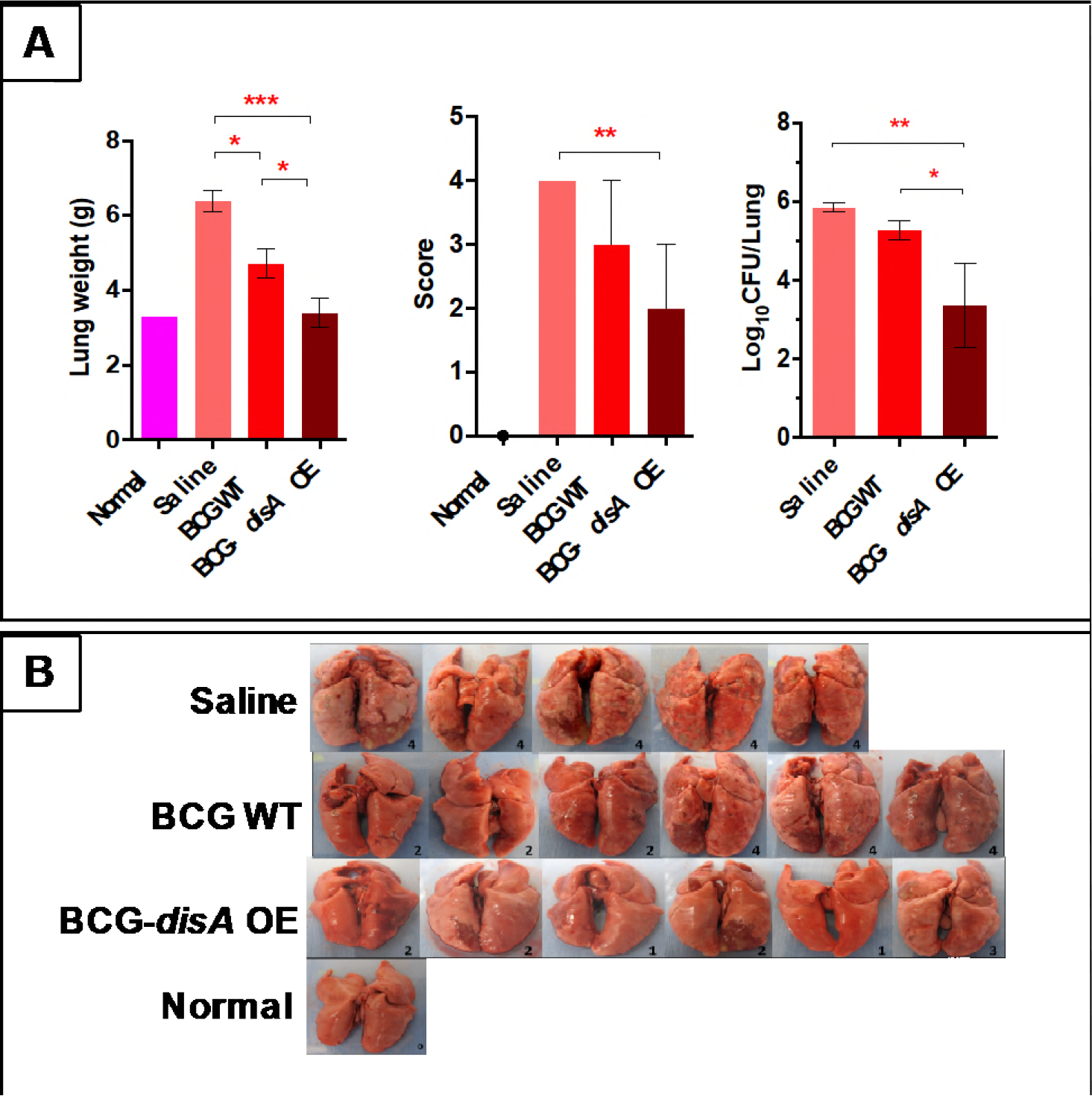
Effect on lung weight, gross lung pathology scores, bacterial burden, and lung gross-morphological features in guinea pigs 18 weeks post-challenge with *M. tuberculosis* H37Rv following vaccination with BCG-WT or BCG-*disA*-OE. A) Lung weights, gross pathology scores, and *M. tuberculosis* bacterial burden at 18 weeks post- *M. tuberculosis* challenge. Two guinea pigs in BCG-*disA*-OE group had lung CFU counts below the limit of our detection. B) Images of lungs at necropsy. ***P < 0.001, **, P< 0.01; *, P<0.05; Non-parametric Man Whitney Test.

Additionally, vaccination with BCG-*disA*-OE effectively controlled the hematogenous spread of *M.tb* to the spleen as evident from significant reduction in spleen weights, spleen pathology scores, and spleen bacterial burdens when compared with the sham-immunized animals at both 14 and 18 week post-infection (**Supplementary Figure 5** and **Figure 4**, respectively). While spleen pathology scores were comparable between animals vaccinated with wild-type and *disA* overexpressing BCG at 14-weeks post-challenge (**Supplementary Fig 5**), by 18 weeks after challenge significantly spleen lower pathology scores was observed in the latter group (**Figure 4**). In addition, there was a trend towards lower spleen CFU in animals vaccinated with *disA* overexpressing strain compared to BCG, especially when examined at 18-weeks post-challenge (**Supplementary Figure 4**). Indeed, at 18 weeks post-challenge, three out of six guinea pigs in BCG-*disA*-OE group had spleen CFU below the limits of detection. These findings clearly indicate that administration of BCG-*disA*-OE effectively controlled *M.tb* replication in the lungs and its dissemination to the spleen.

**Figure 4.**
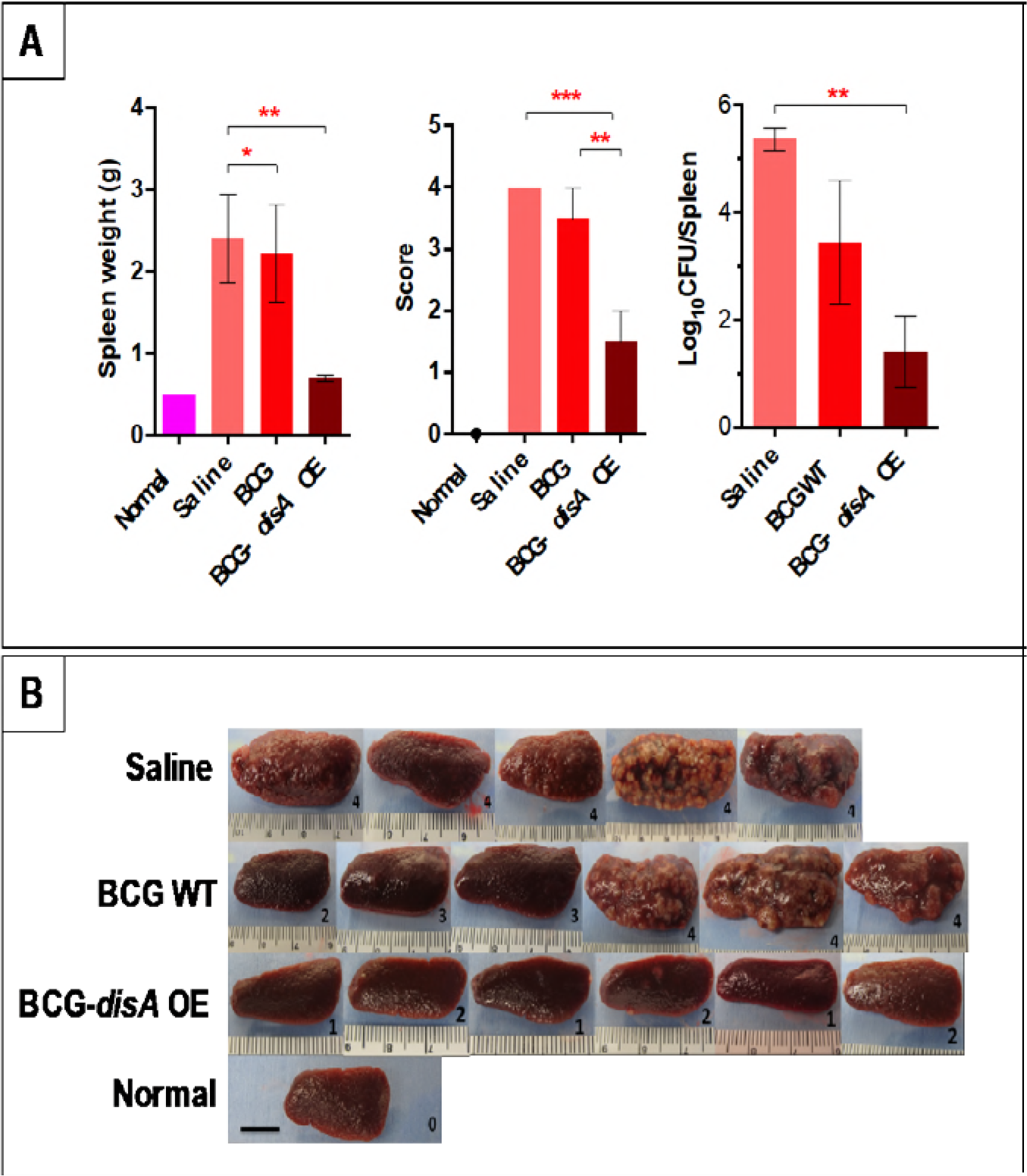
Effect on spleen weight, gross spleen pathology scores, bacterial burden, and spleen gross-morphological features in guinea pigs 18 weeks post-challenge with *M. tuberculosis* H37Rv following vaccination with BCG-WT or BCG-*disA*-OE. A) Spleen weights, gross pathology scores, and *M. tuberculosis* CFU counts at 18 weeks post-*M. tuberculosis* challenge. Three guinea pigs in BCG-*disA-*OE group had spleen CFU counts below the limit of our detection. B) Images of spleens at necropsy. Scale bars indicate 1 cm. The number in the box is gross pathological score. ***P < 0.001, **, P< 0.01; *, P<0.05; Non-parametric Man Whitney Test.

## DISCUSSION

The cytosolic danger sensor STING has emerged as a central Toll-like receptor-independent mediator of host innate immune responses. STING is activated by binding CDNs either secreted by bacteria (such as c-di-AMP or c-di-GMP) or distinct host CDNs (such as cGAMP) generated by host cell receptor following recognition of cytosolic double-stranded DNA [22-24]. As a relatively new class of immunomodulatory molecules, CDNs exhibit strong potential to promote protective immunity and increase vaccine potency through STING-dependent signaling that involve transcription factors IRF-3, IRF-7 and NF-κB [6,10,11]. STING agonists are therefore regarded as promising immune adjuvants for promoting immune responses against tumors and infections. However, the efficacy of small molecule STING agonists as vaccine adjuvants may limited since they are rapidly cleared and may not gain long-lived access to the cytosol for STING activation.

Here we tested the hypothesis that BCG strains overexpressing the STING agonist c-di-AMP might offer sustained intracellular delivery of c-di-AMP and thereby improve the vaccine potential of BCG against TB. We constructed a c-di-AMP–producing recombinant BCG strain by over-expressing the *disA* (MT3692) gene which is identical at the nucleotide level in both *M.tb* and BCG. Since BCG survives and replicates intracellularly for several weeks post-vaccination, over-expression of *disA* under the influence of strong constitutive mycobacterial *hsp60* promoter allows sustained intracellular exposure of the STING agonist in the cytosol of phagocytic cells. By utilizing BCG as the vector for STING agonist delivery, BCG-*disA*-OE offers enhanced innate immune activation via the STING pathways in addition to the full antigenic repertoire of BCG.

Complex genomic rearrangements in BCG strains are one of the major contributors of immunological and phenotypic differences that in turn contribute to the variability in the degree of protection offered by BCG [16,25]. A global resurgence of MDR-TB, HIV-TB co-infection has heightened the need for an improved TB vaccine that provides better protection than that of BCG. Additionally, new vaccines must be safe enough to be used in immunocompromised HIV-TB co-infected individuals. Rational modification of live BCG to increase its antigenic repertoire, and with a prior knowledge of attenuation factors and immunity is critically needed [14,17]. While our study did not directly assess the virulence of BCG-*disA*-OE to that of BCG-WT, based on the finding that Mtb-*disA*-OE showed a median time to death in BALB-c mice of 321.5 days compared to 150.5 days for Mtb-WT following an aerosol infection of 3.5 log10 CFU [12], we anticipate that BCG-*disA*-OE is a weaker pathogen than BCG-WT.

Inside the host, both CD4+ and CD8+ T cells are essential for protective immunity against TB. Dendritic cells (DCs) migrating from the alveoli to the draining lymph nodes are crucial for activation of *M.tb* antigen-specific CD4+ and CD8+ T cells and contribute to resistance to *M.tb* [26]. The requirement for a Th1-like T cells response for host immunity against *M.tb* is clear, and recent vaccine development has sought to stimulate both CD4 and CD8 T-cell responses to produce Th1 cytokines [15]. Hence, elicitation of enduring Th1 responses is a desirable feature of candidate TB vaccines. Not only are STING-activating adjuvants known to elicit antigen-specific Th1 responses, but they also elicit Th17 responses and have been shown to confer improved protection against *M.tb* [10,27]. A recent report published while this manuscript was in preparation suggested that the protection efficacy of protein subunit vaccine adjuvanted with small molecule CDNs was durable for up to 12 weeks after *M.tb* challenge in mice, suggesting that a CDN-adjuvanted vaccine can reduce TB progression in mice through T cell-dependent mechanisms [27].

Guinea pigs are highly susceptible to *M.tb* infection, and the model provides an important pre-clinical evaluation of potential vaccine candidates [28]. Immunization with a single dose of BCG-*disA*-OE resulted in a marked reduction in the gross pathology and bacterial loads in both lungs and spleens of *M.tb* challenged animals when compared to sham treatment or vaccination with BCG alone. Thus our studies highlight the improved potential of BCG-*disA*-OE over BCG to impart protection against *M.tb* infection and disease dissemination.

This study is an important step forward towards implementing CDN-based STING agonists as novel vaccine adjuvants into vaccine strategies for TB. Our results provide proof-of-concept data for utilizing BCG as the vector for producing STING agonist(s) naturally from within the intracellular compartment in a sustained fashion. Our results suggest that STING agonist overexpressing BCG may offer greater efficacy as TB vaccine than BCG alone and that this approach may have utility as an immunotherapeutic tool against other diseases including cancer.

## MATERIALS AND METHODS

**Animals:** All procedures involving live animals were performed in agreement with the protocols approved by the Institutional Animal Care and Use Committee at the Johns Hopkins University School of Medicine. Pathogen-free female outbred guinea pigs (300 g) and C57BL/6 mice were purchased from Charles River Laboratories (North Wilmington, Mass.). Uninfected guinea pigs were housed under pathogen-free conditions at BSL3 animal facility without cross-ventilation. C57BL/6J mice were housed in BSL2 animal facility at the School of Medicine, Johns Hopkins University. Animals were given free access to water and standard mouse or guinea pig chow, respectively. The general behavior and appearance were monitored by veterinary specialists.

**Bacterial strains and cell culture:** Details of all bacterial strains are provided in **Supplementary Table 1**. BCG Pasteur was the kind gift of Frank Collins from the FDA, and M. tuberculosis H37Rv and CDC1551 were obtained from ATCC. Frozen vials of bacterial strains were revived and subsequently sub-cultured in 7H9 Middlebrook liquid medium (B271310, Fisher Scientific) supplemented with oleic acid-albumin-dextrose-catalase (OADC) (B11886, Fisher Scientific), 0.5% glycerol (G5516, Sigma) and 0.05% Tween-80 (BP338, Fisher Scientific) in BSL3 facility. Murine bone marrow was isolated from 4-6 weeks old female C57BL/6J mice. Approximately 10^8^ cells were stored in cryopreservation media made of 10% DMSO (D2650, Sigma) in heat inactivated FBS (10082-147, Fisher Scientific) overnight at −80°C followed by transfer to deep cryopreservation in liquid nitrogen. For differentiation of bone marrow cells into primary macrophages, bone marrow cells were differentiated for 7 days in presence of RPMI-Glutamax (61870-036, Fisher Scientific) supplemented with 10% heat inactivated fetal bovine serum and antibiotics (Penicillin-Streptomycin solution) (15140-122, Fisher Scientific) and 30% (vol/vol) L929 conditioned media. Mouse fibroblast L929 cells (ATCC^®^ CCL-1™) were maintained in RPMI-1640 medium supplemented with 10% FBS and antibiotics.

**Overexpression of MT3692 in BCG Pasteur:** High molecular weight genomic DNA was isolated using CTAB method from log phase cultures of *M.tb*-CDC 1551. Using gene-specific primers (**Supplementary Table 2**), pSD5hsp60.MT3692 (F) and pSD5hsp60.MT3692 (R), the *disA* (MT3692) gene of *M.tb*, was PCR amplified from *M.tb-*derived genomic DNA. Gene amplicons were cloned into mycobacterial shuttle vector pSD5-hsp60 (**Supplementary Table 1**) at the NdeI and MluI restriction sites. The clone (pSD5-hsp60-MT3692) was confirmed by insert release and sequence analyses. The construct pSD5-hsp60-MT3692 was subsequently used to transform BCG. Recombinants clones (BCG-*disA*-OE) selected against kanamycin (25 µg/mL) and further conformed using colony PCR using kanamycin-specific primers (**Supplementary Table 2**). MT3692 overexpression phenotype of BCG-*disA*-OE strain was further confirmed by mRNA expression using quantitative real time PCR (qPCR).

**Quantitative real-time PCR (qPCR):** Late log phase BCG culture pellets were bead beaten using Zirconium beads (KT03961-1-102.BK, Berlin Technologies) before performing total RNA isolation. Trizol reagent (15596026, Fisher Scientific) was used for RNA isolation from both bacterial and mammalian cells. Quantification of mRNA, amplification, and quantification of cDNA was carried out using SYBR Fast green double stranded DNA binding dye (4085612, Applied Biosystems, USA) and ABI StepOnePlus Real Time PCR System (Applied Biosystems, USA). Amplification of *sigH* and mouse beta actin was used as internal controls for BCG and mouse BMDMs respectively. Melt curve analyses confirmed formation of desired and specific PCR product. Experiments were performed in triplicate using three independent biological samples and results were analyzed and presented using 2^-∆∆CT^ method. Details of NCBI gene identifiers and primer sequences are mentioned (**Supplementary Table 2**).

**Macrophage infection, IRF3 activation assay and cytokine ELISAs:** Infection assays were performed in resting mouse BMDMs in 24 well plates in triplicates. Briefly, early log-phase cultures of BCG strains were washed, diluted appropriately to pre-defined concentrations using macrophage infection media (DMEM with 10% FBS) and deposited on the monolayer of cells at a precalibrated MOI (1:20). Infection was allowed continue for 5 h, following which uninfected extracellular bacilli were removed by repeated washing using Dulbecco’s PBS (DPBS). This time point was considered 0 h and the cells were incubated for desired number of hours till the end points were met. To access accurate bacterial counts of infection and internalized bacterial numbers, serial dilutions of the bacterial suspension and 0.025% SDS lysed macrophage were plated on 7H9 plates. RAW-Blue ISG (InvivoGen) reporter cells, derived from the murine RAW 264.7 macrophage cell line by stably integrating an interferon regulatory factor (IRF)-inducible secreted embryonic alkaline phosphatase (SEAP) reporter construct were used for IRF activation assay. The cells were infected with wild-type and BCG*-disA*-OE strains. Cells were incubated for 18 h in fresh macrophage infection media (DMEM with 10% FBS), and culture supernatants were harvested for determination of IRF activation by a SEAP colorimetric assay using QUANTI-Blue reagent (InvivoGen). Following macrophage infection, culture supernatants were isolated, filtered and immediately frozen at −80 for cytokine quantification. Mouse DuoSet ELISA kits for TNF (DY410), IL-6 (DY206-05) and IL1β (DY201-05) were used for cytokine quantification. The absolute concentrations were determined by referring to a standard curve and expressed as pg/mL. For quantification experiments were performed in triplicates.

**Guinea pig immunization and determination of protective efficacy:** To test the prophylactic potential of BCG-*disA*-OE as a vaccine candidate, guinea pigs (n=12 per group) were immunized intradermally using 10^5^ cfu/100 µl of wild-type parental BCG Pasteur or BCG-*disA*-OE strains. Guinea pigs were sham immunized with saline. Animals were challenged with ~100 cfu of *M.tb* H37Rv strain by the aerosol route 6 weeks (**Supplementary Figure 1**) after primary immunization. Lungs from one set of infected animals were harvested, and homogenates were plated on day 1 after to check for established implantation. Infected animals from each group were euthanized 14 and 18 weeks later to determine the protective efficacy of the BCG-*disA*-OE. Gross-pathological features and bacillary burden in lungs and spleen of sham and BCG-immunized guinea pigs after *M.tb* challenge were measured as described previously [29]. **Supplementary Figure 3** shows the details of experimental plan and different animal groups used in the study and **Supplementary Table 1** depicts the bacterial strains and plasmid used in the study.

**Statistical analyses:** Fold-expression (qRT-PCR and ELISA) were represented as mean value ± standard error mean (SEM). Differences between individual test groups were analyzed using by applying unpaired Student’s t-test. All statistics was performed using GraphPad Prism Version 5.01. P values < 0.05 were considered statistically significant.

## ACKNOWLEDGEMENT

The authors thank Dr. Geetha Srikrishna for help with manuscript writing and editing. This work was funded by National Institutes of Health grant R01-AI037856 and HL133190 to W. Bishai.

## SUPPLEMENTARY FI GURES AND TABLES

**Supplementary Figure 1.**
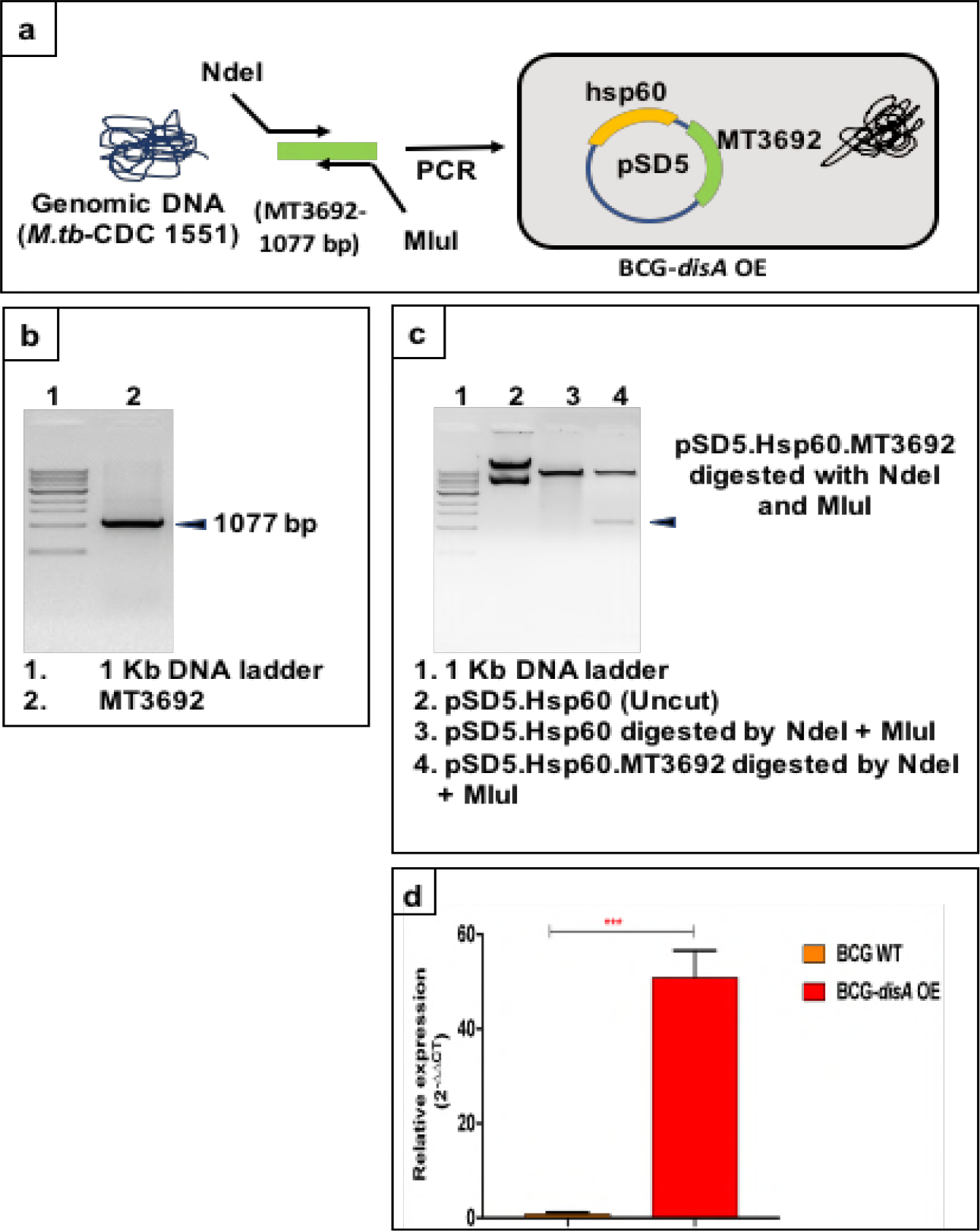
Generation of the BCG-*disA*-OE strain and confirmation of c-di-AMP overexpression. **(a)** The MT3692 (*disA*) gene of *M. tuberculosis* was PCR471 amplified from *M. tuberculosis* CDC 1551 genomic DNA using gene-specific cloning primers. **(b)** The amplicons were cloned into the mycobacterial shuttle expression vector pSD5-hsp60 at the NdeI and MluI restriction sites. The construct (PSD5-*hsp60*-MT3692) generation was confirmed using restriction analyses and sequencing. Constructs were used to transform wild-type BCG Pasteur strain and recombinant clones were selected against Kanamycin (25 μg/mL). **(c)** Differential expression of *disA* in wild-type and BCG-*disA*-OE strains. Gene expression was measured in total RNA isolated from the late log phase cultures using SYBR based quantitative real-time PCR. The graphical data points represent the mean of 3 independent experiments ± standard error mean (SEM). *M. tuberculosis sigA* (Rv2703) was used an internal control. Data analysis was performed using 2^−ΔΔCT^ method. Student’s t test (***P < 0.0001).

**Supplementary Figure 2.**
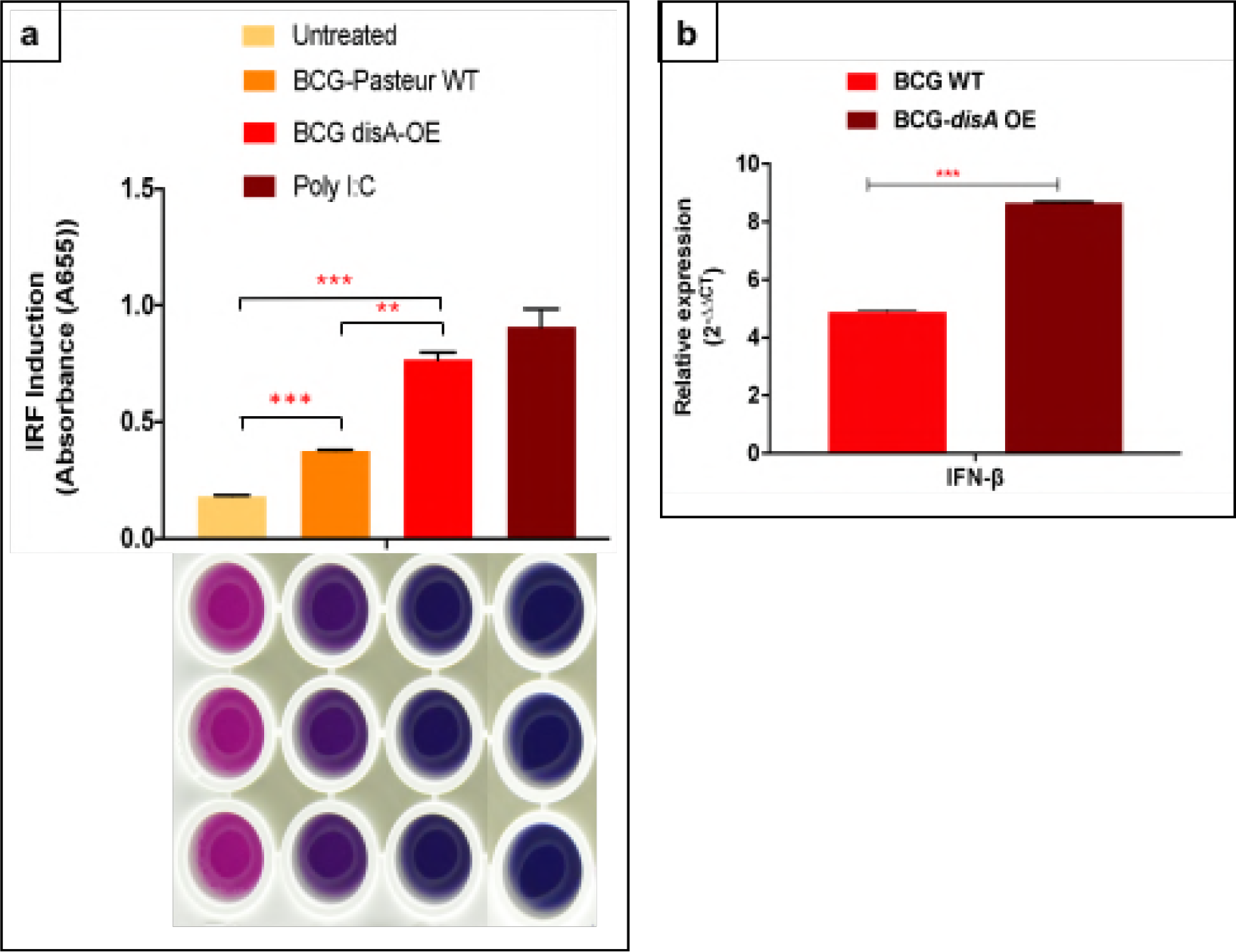
BCG-*disA*-OE overexpressing c-di-AMP gives more potent IRF3 and Type I IFN stimulation that BCG-WT: **(a)** Effect of *disA* overexpression on activation of IRF pathway measured by IRF-SEAP QUANTI Blue reporter assay. RAW-Blue ISG cells were challenged with wild-type and BCG-*disA*-OE strains at an MOI of 1:20 for 5 hours to establish the infection. Uninfected bacteria were washed out using ice-cold DPBS and subsequently incubated for another 18-24 hours. The culture supernatants of infected RAW-Blue ISG cells were assayed for IRF activation. The image below the IRF-activation graph represents QUANTI Blue assay plate and sample wells; treatment parameters for column of wells correspond to those defined for the bars above aligned with the wells. The graphical points represent mean of three independent experiments ± standard error mean (SEM). Student’s t test (***P < 0.0005, **P<0.001). **(b)** Differential expression of IFN*β*: Mouse BMDMs were challenged with wild-type and BCG-*disA*-OE strains at an MOI of 1:20 for 5 hours to establish the infection. Uninfected bacteria were washed using ice-cold DPBS and cells were subsequently incubated for another 6 hours. Expression levels of mRNA was measured using a SYBR green-based quantitative real-time PCR. Basal level of transcript (mRNA) in untreated macrophages was used for data normalization and hence to access relative expression. *β*-actin was used as an internal control. Data analysis was performed using 2^−ΔΔCT^ method. The graphical points represent mean of 3 independent experiments ± standard error mean (SEM). Student’s t test (***P < 0.0005, **P<0.001). MOI (multiplicity of infection).

**Supplementary Figure 3.**
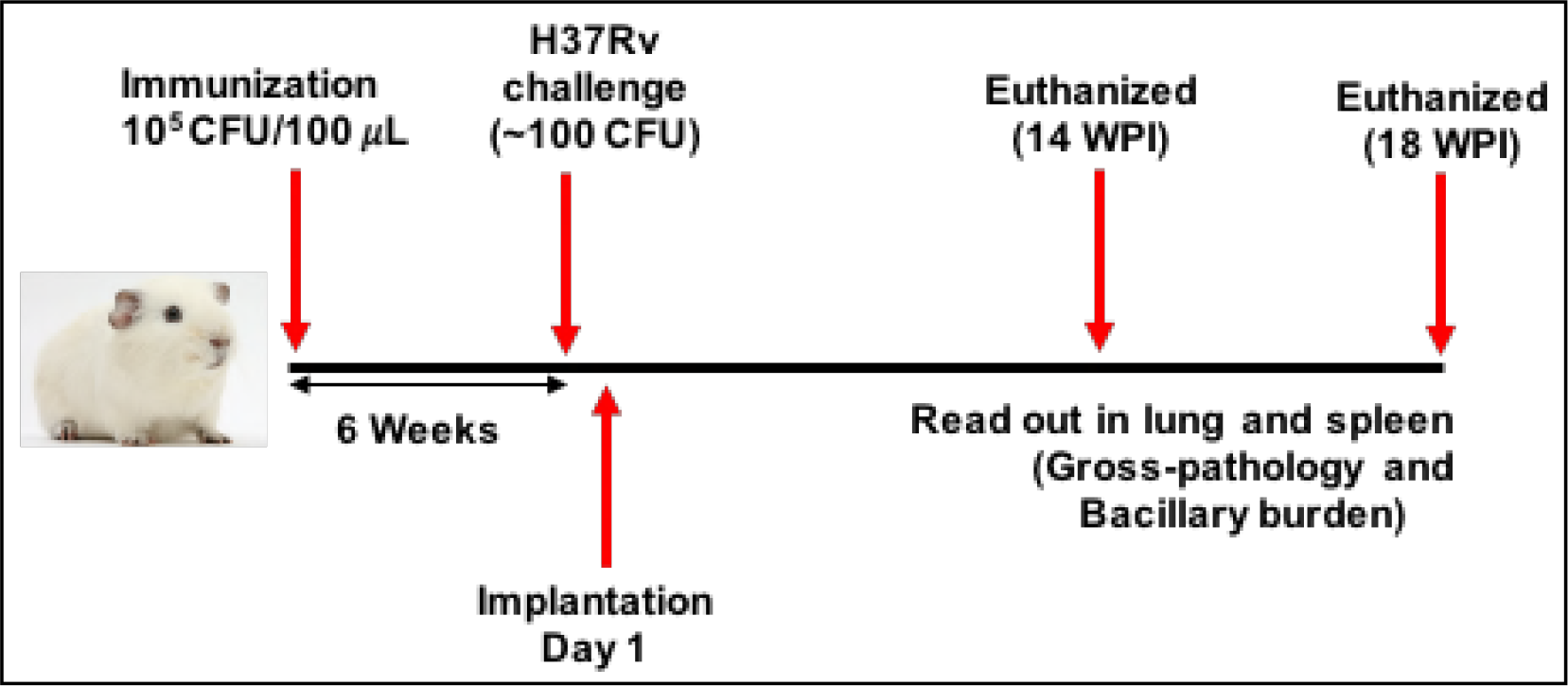
Time line showing the experimental strategy of BCG immunization and challenge in the guinea pig model of vaccination followed by *M.tuberculosis* aerosol infection challenge.

**Supplementary Figure 4.**
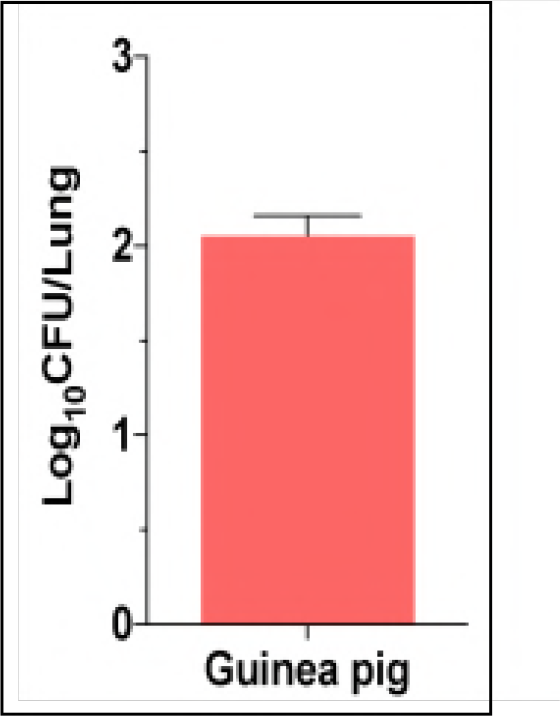
Bacillary load in the guinea pig lung following *M. tuberculosis* challenge. The bacillary load in the lung of guinea pigs (n=3) were determined at day 1 post aerosol challenge. Briefly, animals were euthanized, and lungs were aseptically removed and homogenized in saline. The homogenates were serially diluted and plated in duplicates on 7H11 medium supplemented with appropriate antibiotics. To determine colony forming unit (CFU), Log10 CFU were graphically represented as dot plot, wherein median values ± standard error mean (SEM) are denoted by horizontal line.

**Supplementary Figure 5.**
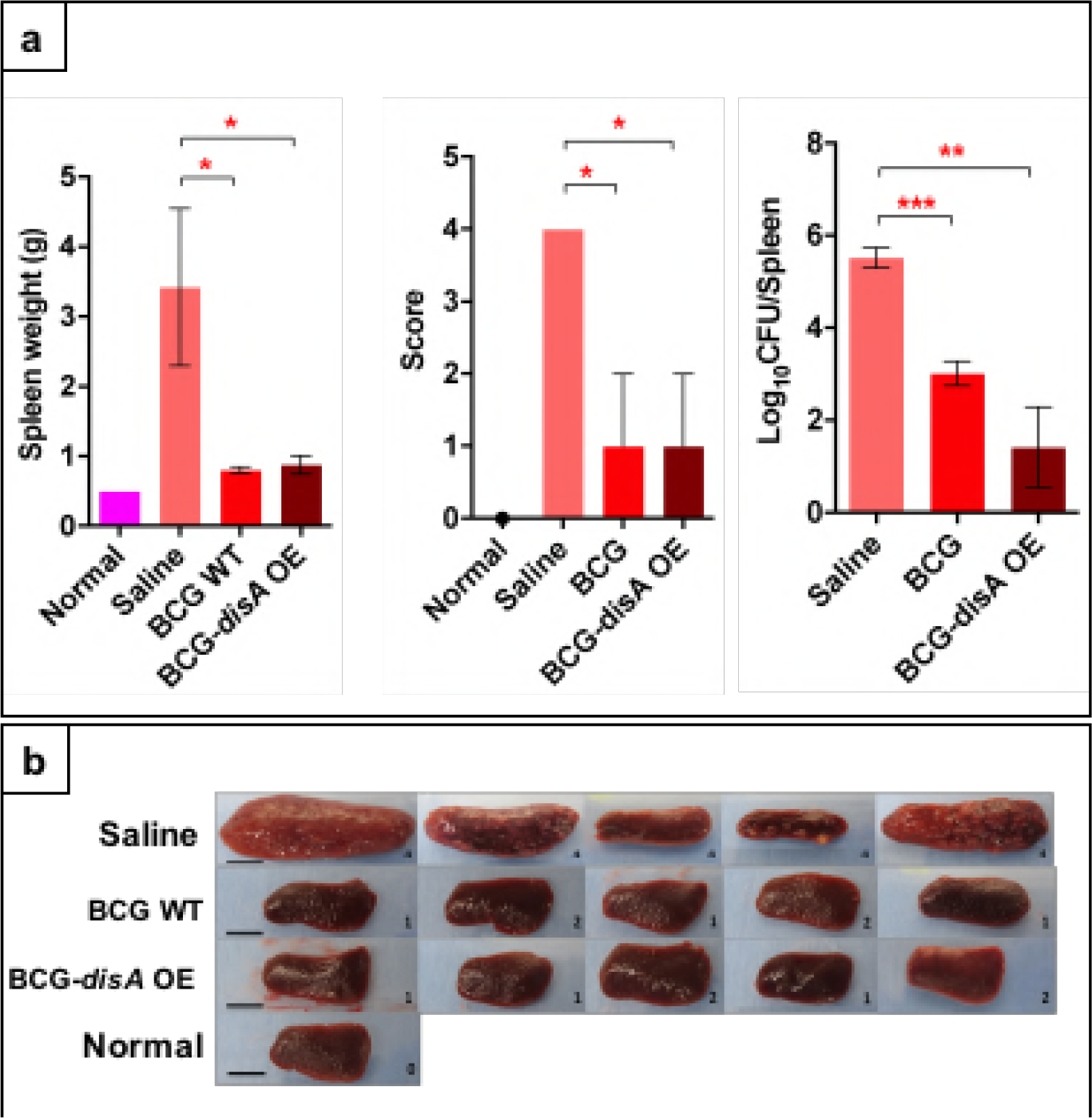
Effect on spleen weight, gross spleen pathology scores, bacterial burden, and spleen gross-morphological features in guinea pigs 14 weeks post-challenge with *M. tuberculosis* H37Rv following vaccination with BCG-WT or BCG-*disA*-OE. A) Spleen weights, gross pathology scores, and *M. tuberculosis* CFU counts at 14 weeks post-*M. tuberculosis* challenge. B) Images of spleens at necropsy. Scale bars indicate 1 cm. The number in the box is gross pathological score. **, P< 0.01; *, P<0.05; Non573 parametric Man Whitney Test.

**Supplementary Table 1:** Bacterial strains and plasmids used in this study.

| Name | Description |
| --- | --- |
| ***M. tuberculosis* strains** |  |
| Mtb-CDC1551 | Wild-type *M. tuberculosis* |
| *Mtb*-H37Rv | Wild-type *M. tuberculosis* |
| ***M. bovis* BCG strains** |  |
| BCG | *M. bovis* BCG Pasteur |
| BCG-*disA*-OE | BCG Pasteur strain overexpressing *disA* (MT3692) of *M.tb* |
| **Plasmids** |  |
| pSD5.hsp60 | Mycobacterial expression plasmid with hsp60 promoter |
| pSD5hsp60.MT3692 | *disA* over-expression plasmid |

**Supplementary Table 2:**
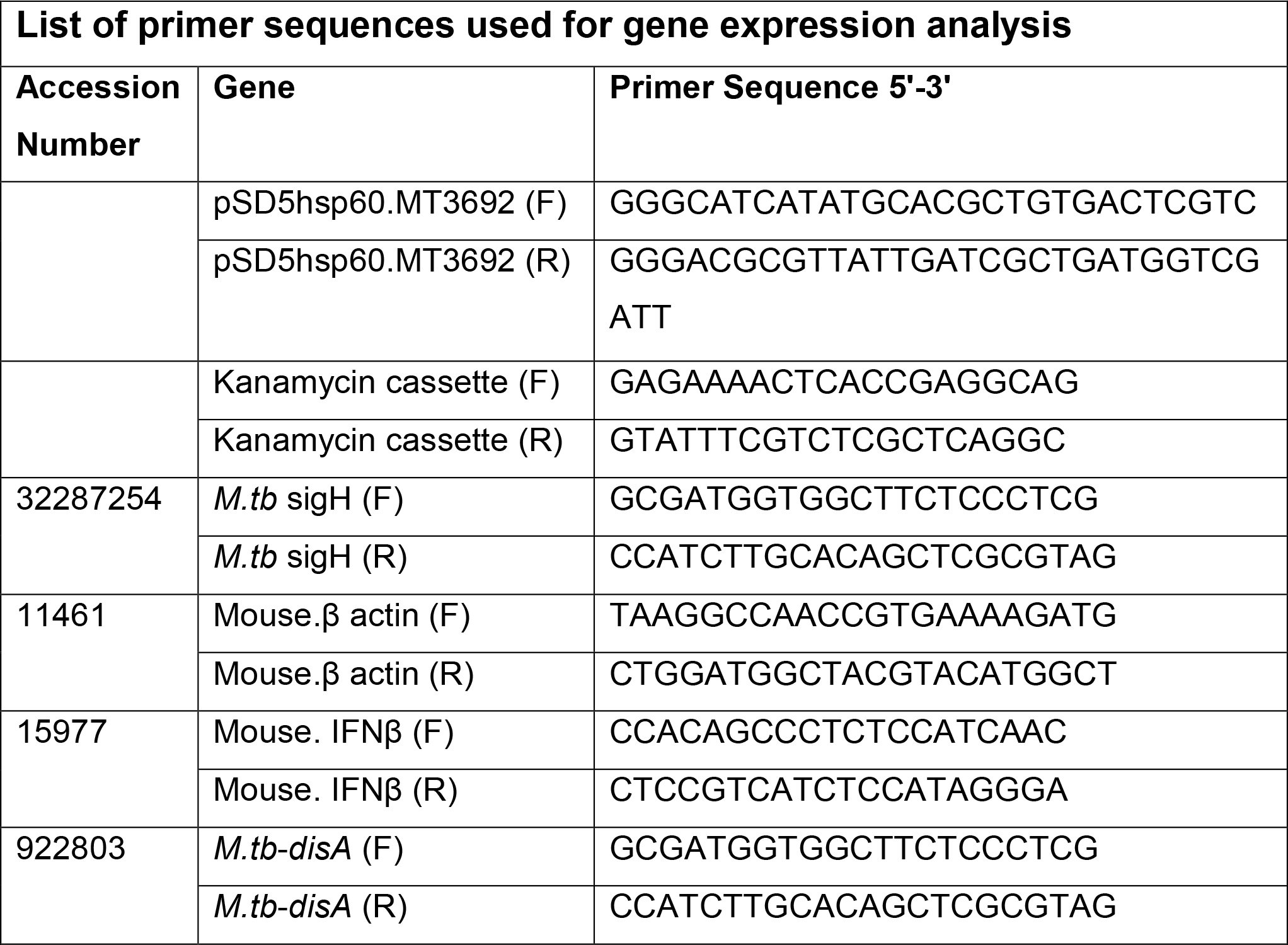
Cloning and PCR primers used in the study.

